# The Third Transmembrane Domain of Synaptophysin Regulates Initial Fusion Pore Dynamics during Ca^2+^-Triggered Exocytosis in Chromaffin cells

**DOI:** 10.1101/2022.01.26.477821

**Authors:** Yu-Tien Hsiao, Meyer B. Jackson

## Abstract

Synaptophysin (syp) is a major secretory vesicle protein comprising four transmembrane domains (TMDs) and a large cytoplasmic C-terminus. The C-terminus of syp has been shown to regulate exocytosis, vesicle cycling, and synaptic plasticity, but the roles of its TMDs remain unclear. SNARE TMDs line initial fusion pores, and structural work along with sequence analysis suggest that TMD III of syp may play a similar role. To test this hypothesis, we expressed TMD III tryptophan mutants in chromaffin cells from mice lacking both syp and its homolog synaptogyrin, and used amperometry to evaluate fusion pores. In contrast to SNARE TMDs, tryptophan substitutions in syp TMD III had no effect on the flux through initial fusion pores. However, these mutants increased the fraction of kiss-and-run events and decreased the initial fusion pore lifetime. Thus, syp TMD III does not line the initial fusion pore, but interacts with it to influence its stability and choice of release mode.

## Introduction

Calcium-triggered exocytosis depends on the coordinated activity of a key set of proteins that drive the fusion of the vesicle and plasma membranes. This process leads to the release of chemical signals throughout the nervous and endocrine systems. Exocytosis begins with the opening of a fusion pore that allows a small trickle of release. The initial fusion pore is formed by the transmembrane domains (TMDs) from two *S*oluble *N*-ethylmaleimide-sensitive factor *a*ttachment protein *re*ceptor (SNARE) proteins, syntaxin in the plasma membrane and synaptobrevin (syb) in the vesicle membrane (Han et al., 2004; Han and Jackson, 2005; Chang et al., 2015; Bao et al., 2016; Chang et al., 2017). SNARE proteins also control the dynamics of initial fusion pores (Borisovska et al., 2005; Han and Jackson, 2006; Fang et al., 2008; Ngatchou et al., 2010; Bao et al., 2018; Weiss, 2019), as do SNARE-interacting proteins, such as synaptotagmins (Wang et al., 2001; Wang et al., 2003; Bai et al., 2004; Zhang et al., 2010; Das et al., 2020) and cysteine string protein (Chiang et al., 2014).

Synaptophysin (syp), a member of the tetraspanner family of vesicle membrane proteins (Sudhof et al., 1987) that is abundant in synaptic vesicles (Takamori et al., 2006; Wilhelm et al., 2014) and endocrine dense-core vesicles (DCVs) (Winkler, 1997; Gonzalez-Jamett et al., 2010), also controls fusion pore dynamics (Chang et al., 2021). At synapses there are no obvious defects in morphology and synaptic transmission in the absence of syp (Eshkind and Leube, 1995; McMahon et al., 1996). However, double knockout (DKO) mice lacking both syp and another tetraspanner, synaptogyrin (syg), display impaired short-term and long-term synaptic plasticity (Janz et al., 1999). Syp KO chromaffin cells show very little reduction in DCV release, but secretion is reduced two-fold in DKO chromaffin cells (Chang et al., 2021). The apparent tolerance to loss of syp and syg may reflect redundancy of function of other tetraspanners.

Tetraspanners harbor four TMDs and a large cytoplasmic C-terminus. The C-terminus of syp influences the time course of catecholamine efflux during DCV exocytosis (Gonzalez-Jamett et al., 2010), controls the dilation and closure of fusion pores (Chang et al., 2021), and regulates the kinetics of synaptic vesicle endocytosis (Kwon and Chapman, 2011). However, the roles of the TMDs of syp remain unclear. Purified syp forms hexameric channel-like structures (Arthur and Stowell, 2007) and produces channels in lipid membranes (Thomas et al., 1988). Syp has a similar overall TMD topology to connexins, the gap-junction channel proteins (Betz, 1990; Leube, 1995; Adams et al., 2015). The third TMD of connexin lines the gap junction pore and is characterized by charged, polar, or small residues alternating with large hydrophobic residues. The third TMD of syp displays a similar pattern (Betz, 1990). Weak sequence homology between syp TMD III and connexin TMD III have been noted (Leube, 1995; Arthur and Stowell, 2007). Furthermore, in a complex containing 6 copies of syp and 12 copies of syb, TMDs III and IV of syp partner with the TMD of syb to line a central aqueous cavity (Adams et al., 2015). In this complex the residues of syb that face inward include those that alter catecholamine flux through endocrine fusion pores (Chang et al., 2015; Chang et al., 2017) and glutamate flux through synaptic fusion pores (Chiang et al., 2018). These studies raise important questions about the role of tetraspanner proteins in fusion pores. If a cavity like that seen in the syb-syp structure forms at the onset of fusion, then syp TMD III would be a pore liner and influence pore flux in a manner similar to that seen with the SNARE TMDs.

To address these questions, we tested syp TMD III tryptophan mutants in chromaffin cells, and used amperometry to assess their impact on fusion pores formed during calcium-triggered DCV exocytosis. DKO cells provide an advantageous background for these studies because they lack the two major endogenous chromaffin cell tetraspanners that could obscure the effects of exogenous mutants. We found that wild type and TMD III mutants support exocytosis in DKO chromaffin cells. The actions of these mutants indicate that syp TMD III regulates the dynamics of initial fusion pores and the mode of release, without influencing late-stage fusion pores. Tryptophan substitutions throughout TMD III failed to alter the catecholamine flux through the initial fusion pore, indicating that this domain is not part of the pore lining. Thus, the pore-like syp-syb complex (Adams et al., 2015) does not represent the initial fusion pore of Ca^2+^-triggered exocytosis. Although it is not a pore liner, syp TMD III does play a role in determining initial fusion pore stability, and in controlling whether it dilates or closes.

## Results

### Syp TMD III mutants support secretion

To investigate the role of syp TMD III we replaced 12 different residues of TMD III with tryptophan and expressed these mutants in DKO chromaffin cells. To confirm that exogenous wild type syp and tryptophan mutants target to DCVs, we co-expressed a neuropeptide Y fusion construct, NPY-DsRed, to mark DCVs in DKO cells. We then evaluated colocalization with syp by immunocytochemistry. Wild type syp and selected mutants (green) colocalized with NPY-DsRed (red) (Fig. 1A), and had no significant differences between their Pearson’s correlation coefficients (Fig. 1B). This confirms the presence of syp on DCVs (Winkler, 1997; Gonzalez-Jamett et al., 2010; Chang et al., 2021), and suggests that our mutations in syp do not affect their targeting to DCVs. Since syp associates with syb and syb targets to DCVs (Chang et al., 2015), we also checked the colocalization of syp and syb. As expected, wild type and mutant syp (red) colocalized with syb (green) (Fig. 1C) and the Pearson’s coefficient was comparable between wild type syp and mutants (Fig. 1D). Note that Pearson’s coefficient with syb in Fig. 1D was higher than that with NPY in Fig. 1B, suggesting that syp targets some structures that are not DCVs, such as small clear vesicles. These results suggest that exogenous wild type and mutant syp efficiently targets DCVs.

**Figure 1.**
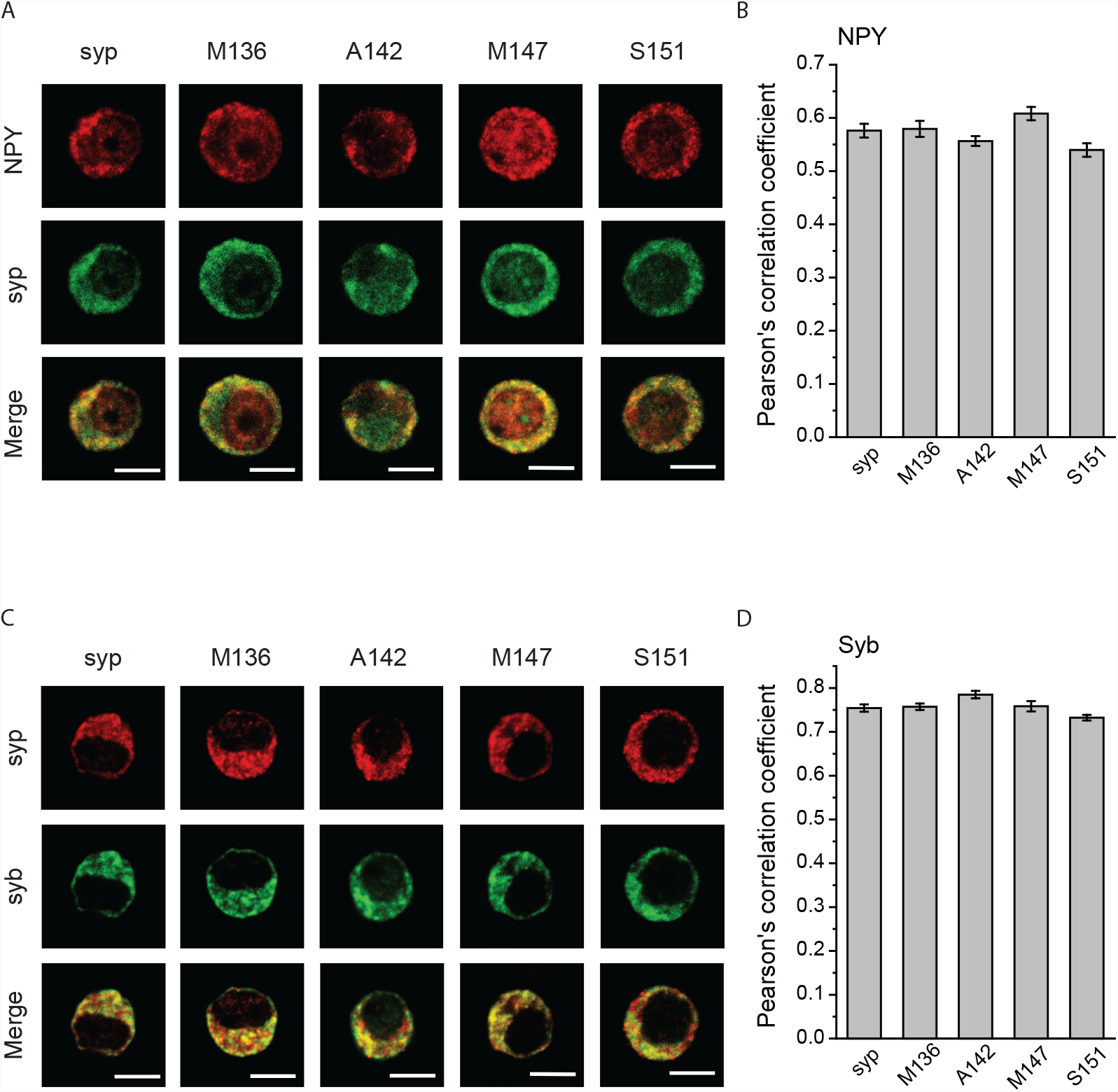
Vesicle targeting by wild type and mutant syp. **A**. Immunostaining images of NPY-DsRed (red) and syp (green) in DKO chromaffin cells co-transfected with NPY-DsRed and wild type syp or with the indicated syp tryptophan mutants. **B**. Pearson’s correlation coefficient for colocalization of syp and NPY. **C**. Immunostaining of syp (red) and syb (green) in DKO chromaffin cells transfected with wild type syp or tryptophan mutants. **D**. Pearson’s correlation coefficient for colocalization of syp and syb. N = 18-28 cells. Scale bar: 5 μm.

We then used amperometry to test the impact of these mutants on exocytosis. Fig. 2A presents amperometry recordings from control DKO cells, and DKO cells expressing wild type syp, syg, and 3 mutants. Upon depolarization by application of a high-KCl solution, untransfected DKO cells and DKO cells expressing wild type syp or syg produce amperometric spikes at similar frequencies, which are roughly half that seen in wild type chromaffin cells (Chang et al., 2021). Thus, although DKO cells lack the two major tetraspanners, they do not provide a secretion-free background, possibly due to redundancy of function with other tetraspanners. Spike frequency did not have a statistically significant dependence on syp form (by ANOVA), but tryptophan substitution in the N-terminal part of TMD III tended to increase frequency while substitutions that reduced frequency were only found in the C-terminus (Fig. 2B). Cumulative plots of spike number versus time provide another view of spike frequency by focusing on the 10-70% rising-slope (Fig. 2C). This analysis indicated some variation depending on position of tryptophan substitution: most mutants toward the N-terminus produced a faster rising-slope, while three mutants toward the C-terminus (L149, V150, and S151) reduced the rising-slope (Fig. 2D). One mutant near the C-terminus (S152) produced a small increase in rising-slope. Note that spike frequency (Fig. 2B) provides a statistically more conservative analysis based on cell-mean, whereas rising-slope (Fig. 2D) combines the total number of spikes without using the number of cells. Thus, the error in the rising-slope is smaller because it depends on the much larger event number. The appropriateness of event-based and cell-based statistical analysis of amperometry data depends on the contribution of variation between cells to the total variance (Colliver et al., 2000; Chang et al., 2021). With this qualification, the results indicate that these tryptophan mutants exert neither a large dominant negative action nor a rescue action. However, the parallel trends in spike frequency and rising-slope, and the clustering of similar actions, suggest that tryptophan substitution toward the N-terminus can favor secretion, while substitutions in the C-terminus can inhibit secretion.

**Figure 2.**
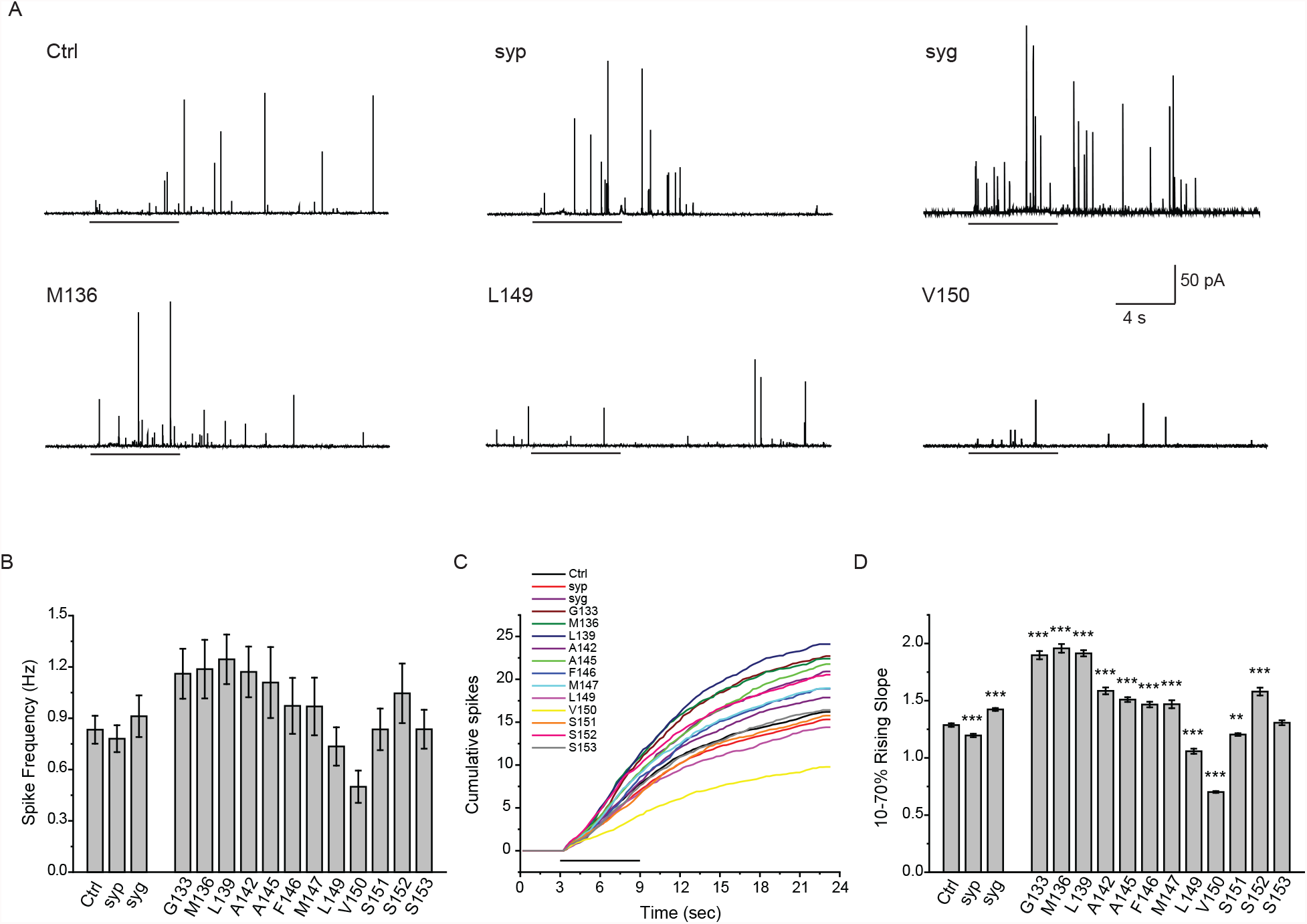
Spike frequency. **A**. Traces of amperometry recordings from DKO cells expressing empty vector (Ctrl), wild type syp, syg, and tryptophan mutants. Black horizontal lines below indicate application of depolarizing, high KCl, solution. **B**. Mean spike frequency averaged over cells. There was no significant difference in all tested groups. **C**. Cumulative spike counts were plotted versus time, with black line representing high KCl application as in **A. D**. 10-70% rising-slopes of cumulative plots determined from linear fits. Data were from 103 cells for Ctrl; 76 cells for wild type syp; 45 cells for syg; 20-41 cells for mutants. ** *p* < 0.01; *** *p* < 0.001, Kruskal-Wallis method followed by *post-hoc* Dunn test.

### Changes in kiss-and-run fraction and duration

Exocytosis begins with the opening of a fusion pore, which is clearly visible in amperometry recordings. Upon opening, a fusion pore can either dilate or close. Dilation leads to full fusion with secretion of the bulk of vesicle content, while closure limits loss to a fraction of total content in a process termed kiss-and-run. We can estimate the fraction of these two outcomes on the basis of peak amplitude (Wang et al., 2006; Chiang et al., 2014; Chang et al., 2021). Events < 4 pA tend to produce a slow steady release and are taken as kiss-and-run events. Fig. 3A displays examples from control chromaffin cells and DKO chromaffin cells expressing two mutants. Based on the amplitudes of these events we used 4 pA as a cutoff to calculate the fraction of kiss-and-run events. Using 6 pA as the cutoff increases this fraction overall while preserving differences between cells expressing different proteins (Chang et al., 2021). Syp KO cells and DKO cells displayed higher fractions of kiss-and-run events, suggesting that syp can influence the choice between kiss-and-run and full fusion (Chang et al., 2021). DKO cells expressing syg showed a comparable fraction to control DKO cells, suggesting that syg has a relatively small influence on release mode. Tryptophan replacement of M136, L139, and A142 in TMD III significantly increased the cell-mean of this fraction (Fig. 3B). Furthermore, two of these three mutants increased the duration of kiss-and-run events compared to other mutants, control DKO cells, and wild type syp and syg (Fig. 3C). This suggests that this part of TMD III plays a role in stabilizing the fusion pore formed during kiss-and-run.

**Figure 3.**
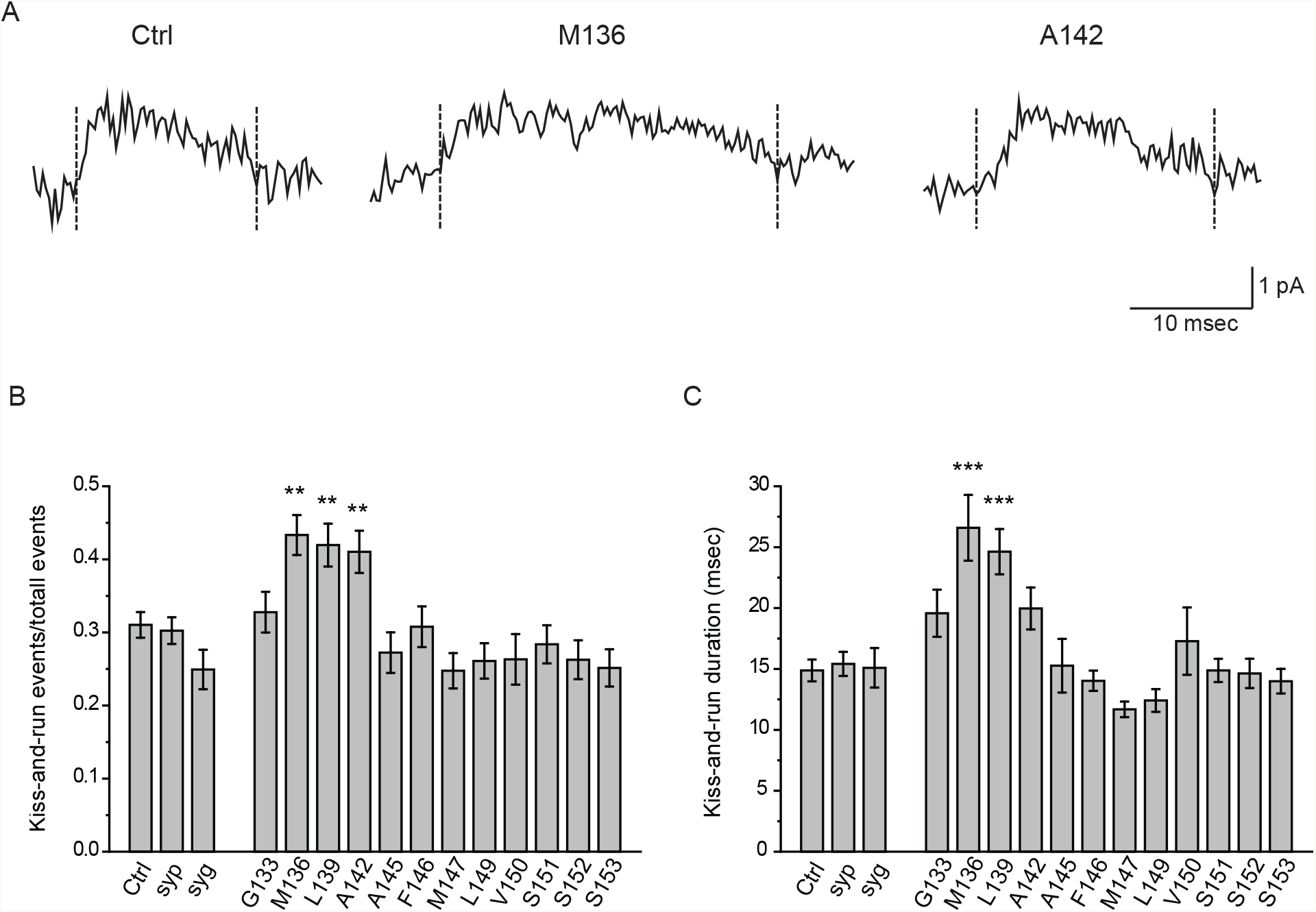
Kiss-and-run events. **A**. Amperometric recordings from Ctrl cells and cells expressing two tryptophan mutants. **B**. Fraction of kiss-and-run events was calculated from events with peak amplitudes of 2-4 pA relative to the total events with peak amplitude ≥ 2 pA. **C**. Kiss-and-run duration in all test groups. Cell numbers as in Figure 2. ** *p* < 0.01; *** *p* < 0.001, One-way ANOVA followed by *post-hoc* Student-Newman-Keuls test.

### Initial fusion pore stability

When the opening of a fusion pore leads to a spike, the small current is referred to as a prespike foot (PSF) (Fig. 4A, gray area) (Chow et al., 1992; Jankowski et al., 1993). To determine the role of TMD III in the stability of these initial fusion pores, we examined PSF in DKO cells expressing tryptophan mutants. The mean PSF duration was determined by fitting a single exponential to the PSF lifetime distribution (Fig. 4B). The resulting values were similar in DKO cells expressing control and wild type syp (Fig. 4C). By contrast, syg shortened the PSF lifetime, suggesting that this protein destabilizes the initial fusion pore. Four tryptophan mutants, M136, F146, L149, and S153, altered the mean PSF duration and in each case the duration was reduced (Fig. 4C). This suggests that these four residues engage in interactions that stabilize the initial fusion pore, and that tryptophan replacement weakens these interactions.

**Figure 4.**
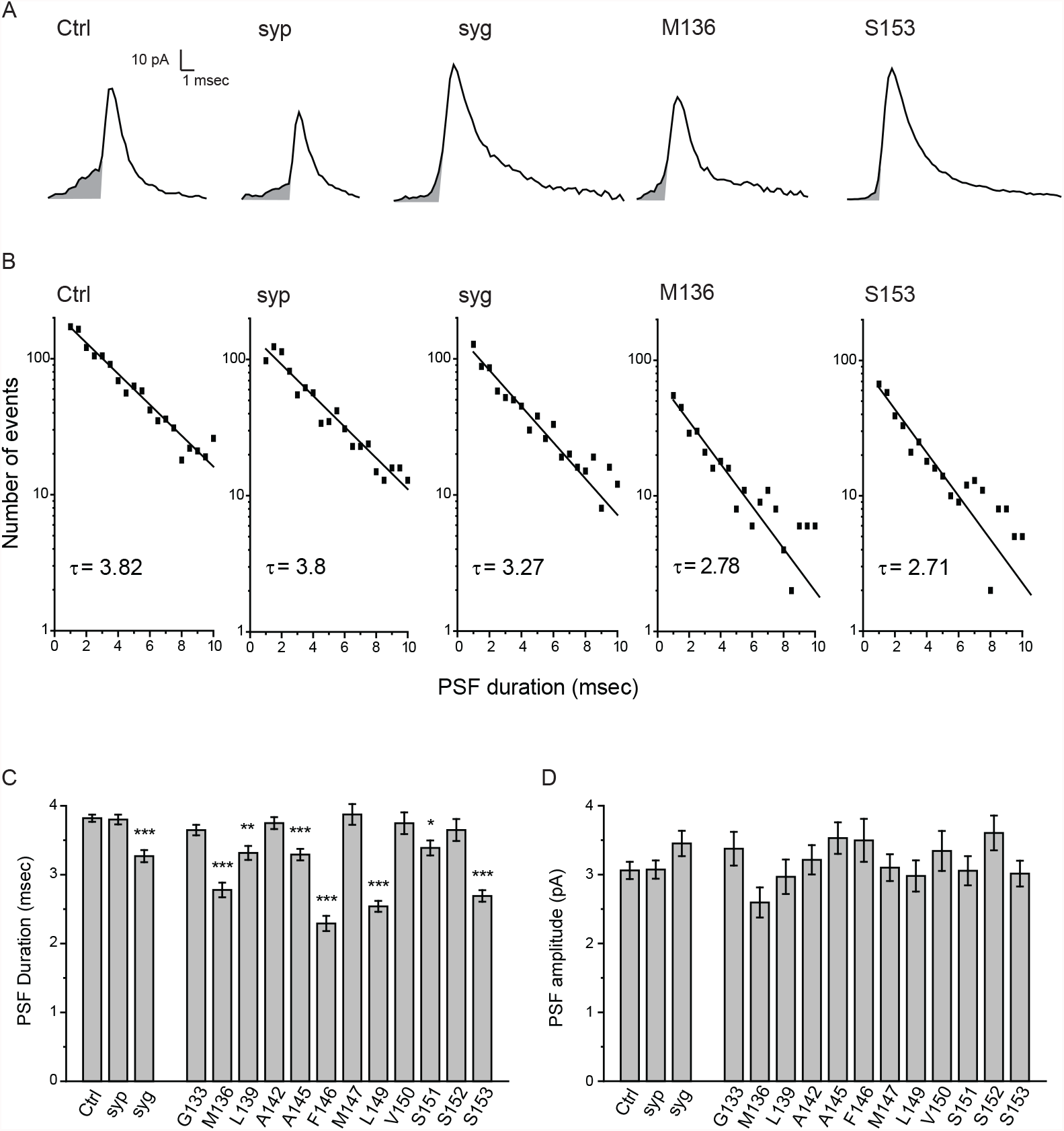
Pre-spike feet. **A**. Amperometric spikes recorded from Ctrl, syp, syg, and two tryptophan mutants, M136 and S153. Pre-spike feet (PSF) are indicated as gray shaded regions. **B**. Lifetime distribution of PSF were fitted with a single exponential decay function to yield τ, the mean PSF duration. **C**. Mean of PSF duration from the fits as in **B. D**. Cell-mean of PSF amplitude (1500 events and 103 cells for Ctrl; 1031 events and 76 cells for syp; 871 events and 45 cells for syg; 249-700 events and 20-41 cells for mutants). * *p* < 0.01; ** *p* < 0.01; *** *p* < 0.001, one-way ANOVA followed by *post-hoc* Student-Newman-Keuls test.

### Initial fusion pores flux

The findings that syp and syb form a complex (Edelmann et al., 1995) with a hexameric pore-like structure (Adams et al., 2015), and that syp TMD III has similar transmembrane topologies to connexin TMD III (Betz, 1990; Leube, 1995), raise the possibility that TMD III serves as a structural component of a fusion pore. Catecholamine flux can be used to probe the size of a pore and tryptophan TMD mutations in SNAREs reduce PSF amplitudes (Han et al., 2004; Chang et al., 2015). We therefore examined the PSF amplitudes in DKO cells expressing syp TMD III mutations. None of the TMD III mutants produced statistically significant changes in PSF amplitude (Fig. 4D). The largest effect with M136 was a reduction of only 16%. By contrast, 3 TMD mutations in syntaxin (Han et al., 2004) and 4 TMD mutations in syb (Chang et al., 2015) produced statistically significant reductions in PSF amplitude. The finding that tryptophan substitutions in TMD III of syp do not influence catecholamine flux through the initial fusion pore indicates that this domain does not reside near the exit pathway taken by catecholamine during its release.

### Spike shape and fusion pore expansion

Fusion pore dilation produces a spike, the shape of which is partly dependent on the speed and extent of this dilation. The C-terminus of syp has been shown to influence these processes (Chang et al., 2021). To determine whether TMD III influences late-stage fusion pores we examined spike shape (Fig. 5A). Spike amplitude (Fig. 5B), 10-90% rising time (Fig. 5C), half-width (Fig. 5D), decay time (Fig. 5E), and whole area (Fig. 5F) were not significantly changed by tryptophan mutations in TMD III. To relate spike shape to late-stage fusion pores, we calculated permeability as the ratio of amperometric current to remaining vesicle catecholamine content (Jackson et al., 2020). As shown previously, permeability rises suddenly as the fusion pore expands, and after reaching a peak decays to a plateau (Fig. 6A). We measured the peak and plateau permeabilities and found they were indistinguishable for the proteins expressed (Fig. 6B and 6C). These results suggest that TMD III has little if any influence on late stage fusion pores.

**Figure 5.**
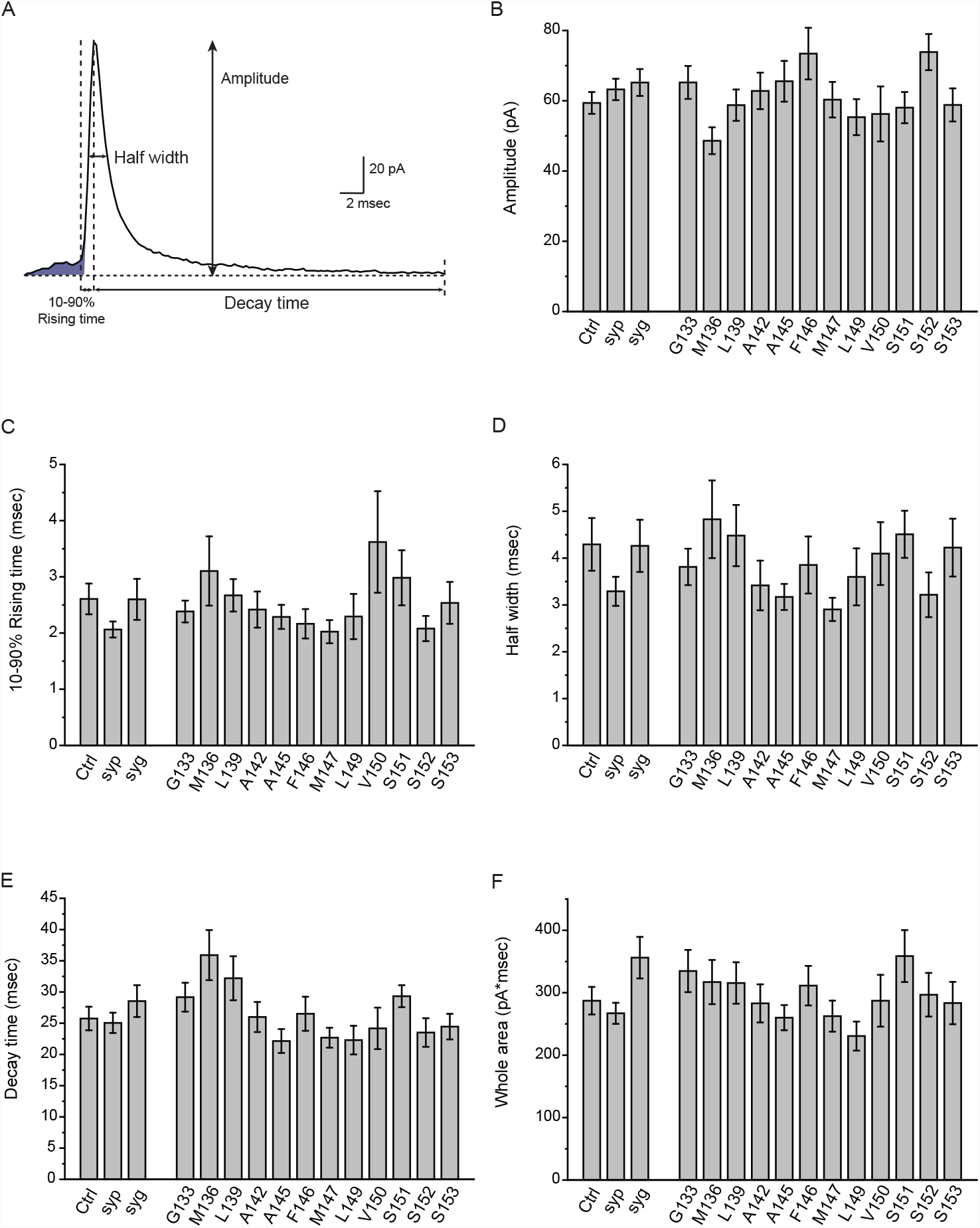
Spike characteristics. **A**. An amperometry spike with PSF shaded and key parameters illustrated. **B-F**. Cell-means of peak spike amplitude (**B**), 10-90% rising time (**C**), half width (**D**), decay time (**E**), and total area (**F**). Values were calculated from spikes with peak amplitude ≥ 20 pA. Cell numbers as in Figure 2.

**Figure 6.**
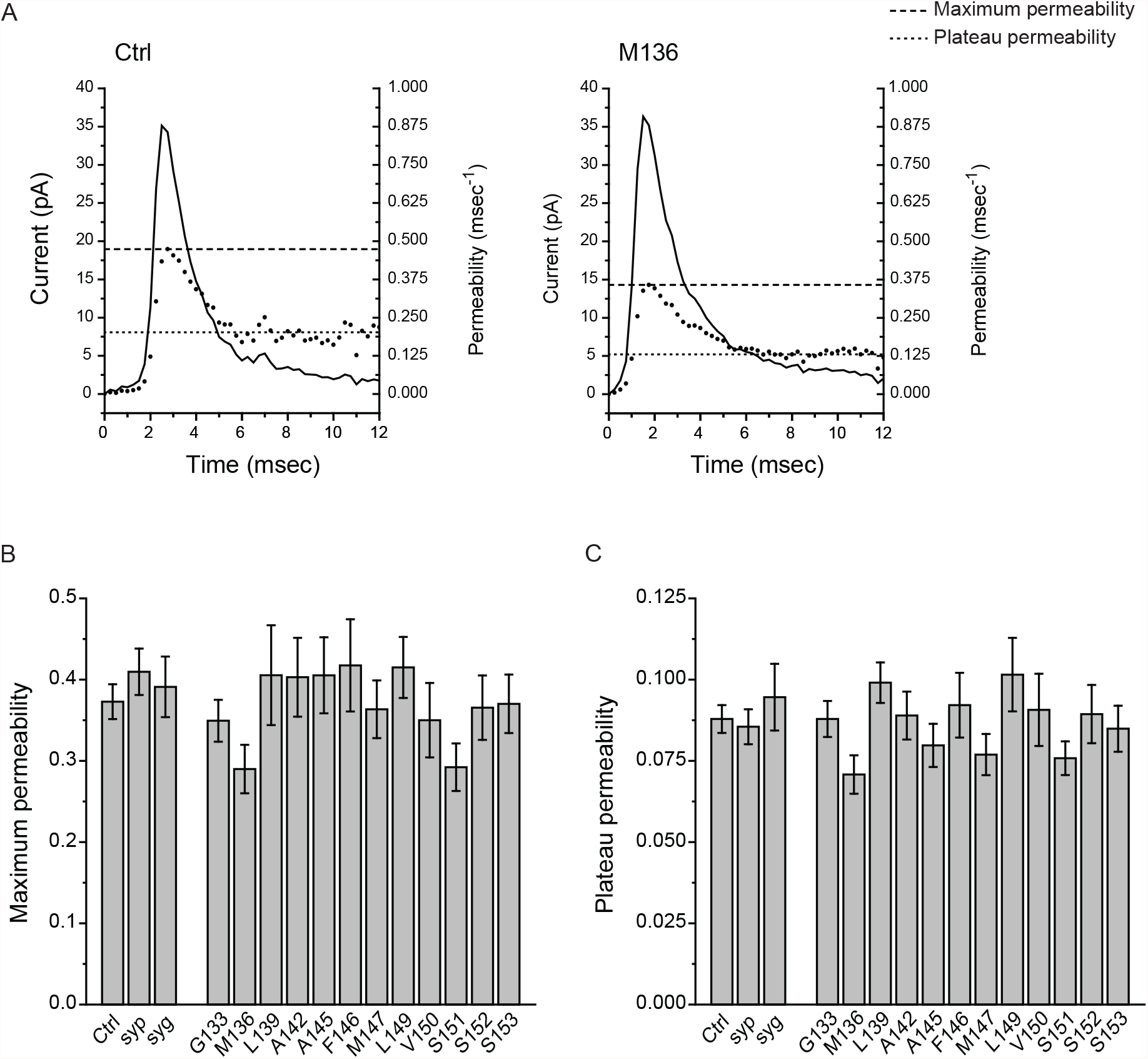
Late stage fusion pores. **A**. Permeability (dotted trace) was calculated by using amperometric currents (solid trace) as described (Jackson et al., 2020). Maximum and plateau permeability are indicated with dashed and dotted lines, respectively. **B**. Peak permeability. **C**. Plateau permeability. Cell-means, with cell numbers as in Figure 2.

## Discussion

Roles for tetraspanners in the fusion of membranes during exocytosis have proven difficult to identify. Overexpressing syp or syg in PC12 cells downregulates human growth hormone release (Sugita et al., 1999). Depleting neuronal syp and syg increased synaptic vesicle release at the calyx of Held synapse (Raja et al., 2019). In *Drosophila* eliminating syg reduced spontaneous synaptic release (Stevens et al., 2012), but *C. elegans* tetraspanner null mutants displayed normal miniature postsynaptic currents (Abraham et al., 2006). Redundant actions of different isoforms make it difficult to evaluate the results of knock-out experiments, and species differences add to the challenge of determining general functions. We recently found that syp protein level and secretion were not well correlated (Chang et al., 2021). Here we showed that exogenous syp and syg do not enhance secretion above that seen in DKO cells (Fig. 2). Experiments with a syp deletion mutant had suggested different functions for the C-terminus and TMDs (Chang et al., 2021). Here we investigated the TMDs in more detail, focusing on TMD III, a domain with a potential role as a fusion pore component. Tryptophan substitutions in TMD III did not alter flux through initial or late-stage fusion pores. Mutations at specific locations did however influence the stability of initial fusion pores and the mode of exocytosis. Thus, although it does not line the fusion pore, TMD III controls functionally important fusion pore transitions in early stages of calcium-triggered exocytosis in endocrine cells.

### Syp TMD III and initial fusion pore composition

Our experiments tested the hypothesis that TMD III of syp lines the initial fusion pore. Syp interacts with syb to form a stoichiometric complex with a channel-like structure (Adams et al., 2015). If this complex represents an initial fusion pore, tryptophan mutants in TMD III that face into the pore lumen would be expected to obstruct the flow of catecholamine and reduce the amplitude of the PSF (Fig. 4D). The failure of these mutations to alter initial fusion pore flux suggests that these residues do not interact with permeating catecholamine. This suggests that despite the homologies with connexin (Betz, 1990; Leube, 1995), it is unlikely that TMD III lines the initial fusion pore. The syp-syb complex characterized by Adams et al (2015) does not appear to represent the initial fusion pore, but other functions for this complex remain a possibility.

### Syp TMD III and fusion pore dynamics

Although it is not a pore liner, TMD III does nevertheless influence initial fusion pores. Changes in PSF lifetime indicate that TMD III influences initial fusion pore stability (Fig. 4). The effect is specific to initial fusion pores as spike shape and late-stage fusion pore permeability were not affected (Fig. 5 and 6). Changes in the proportion and lifetime of kiss-and-run events indicate an additional role in the choice of release mode (Fig. 3). It is notable that mutations with a destabilizing effect on fusion pores leading to spikes (shorter PSF lifetimes), increase the lifetime of kiss-and-run events (longer lifetimes). Thus, these tryptophans pack well in the latter configuration and poorly in the former. This suggests that TMD III is arranged differently in these two fusion pore states. Interestingly, the three residues implicated in the control of release mode are located close to the cytosolic face of the vesicular membrane. Tryptophan within the syb TMD perturbs the surrounding lipid bilayer and alters membrane curvature (Chang and Jackson, 2015). Tryptophan in syp TMD III could similarly impact the stability of a lipidic fusion pore, and thus influence dilating transitions from an initial proteinaceous fusion pore. A previous study indicated that the syp C-terminus influences the choice between full fusion and kiss-and-run (Chang et al., 2021). Thus, both TMD III and the C-terminus of syp can affect the release mode. These results suggest that TMD III is intimately involved in the initial fusion pore. It is not a pore liner itself but may interact with other domains within the initial fusion pore.

### TMD interactions between syp and syb

The most likely interacting partner for syp in the vesicle membrane is syb. Thus, we examined prior results with syb (Chang et al., 2015), and explored the potential role of syp-syb interactions using structural models. Helical wheels (Fig. 7A) illustrate that residues of syb that influence initial pore stability are distributed through most of its TMD. The 7 residues of syp TMD III that influence initial fusion pore stability are also distributed. To explore interactions between these TMDs in greater detail we examined the syb-syp complex of Adams et al. (2015). Although the present results indicate this structure is not directly relevant to initial fusion pores, it nevertheless renders detailed interactions between the TMDs. In this structure 5 of the 7 residues of syp with an effect on PSF lifetime (L139, A145, F146, L149, and S153) face away from the syb TMD and toward syp TMD IV (included in Fig 7B1) or toward syp TMD I (not shown). Two residues of TMD III with an effect on PSF lifetime, M136 and S151, are close to syb. As an alternative we used a model based on Alphafold (Jumper et al., 2021) (Fig. 7B2). Here we see more residues facing away from TMD IV (M136, L139, and F146), and these could interact with another molecule such as syb. S153 probably also faces toward syb because in the helical wheel, L139, F146, and S153 are located on the same face (Fig. 7A). These residues that appear to face toward syb were found to reduce PSF duration (Fig. 4). It should be noted that the syb residues that are potential partners with syp residues are not among those lining the pore (Chang et al., 2015) (purple space fills in Fig. 7B1). These observations raise the possibility that syp TMD III could influence initial fusion pore stability by intermolecular interactions with syb as well as intramolecular interactions with other syp TMDs. Future studies of other syp TMDs may provide more insight into how syp regulates fusion pores, and suggest additional functional interactions with syb and other proteins of the exocytotic apparatus.

**Figure 7.**
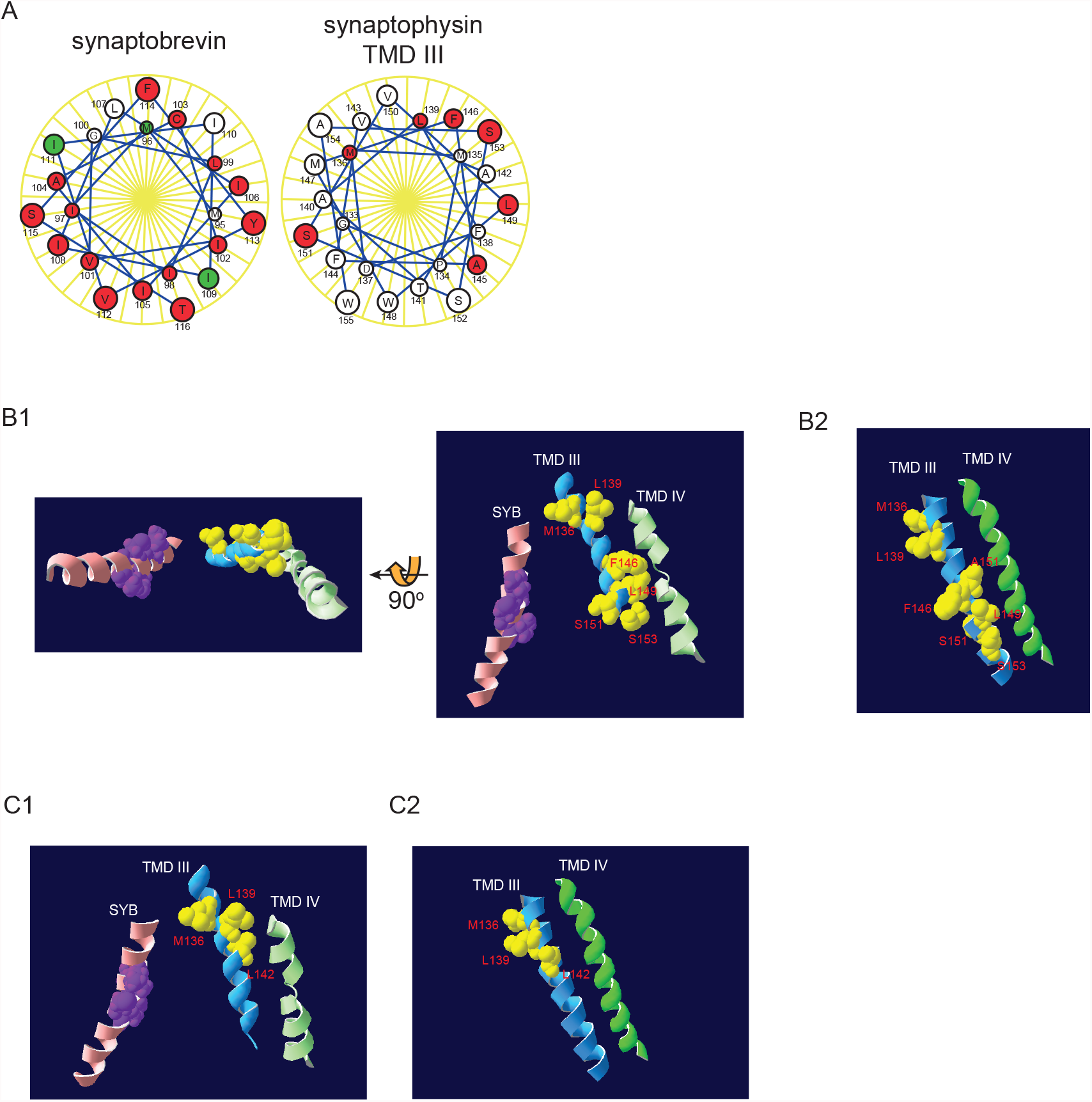
TMDs of syp and syb. **A**. Helical wheels of syb (left) and syp (right). Residues where tryptophan decreased PSF duration are marked in red, and residues where tryptophan increased PSF duration are marked in green. Residues that had no impact or were not tested are white (data for syp are from Fig. 4 and data for syb are from (Chang et al., 2015)). **B**. Structural models of TMDs of syb and syp using a pdb file from (Adams et al., 2015) (**B1**) and the Alphafold database (Jumper et al., 2021) (**B2**)). TMD III (blue ribbon), TMD IV (green ribbon), and syb (pink ribbon) are viewed from outside (**B1**, left) and from within the plane of the membrane (**B1**, right). Residues of syp TMD III that influence the open time of the fusion pore are highlighted as yellow fills. Syb pore-lining residues are purple fills (Chang et al., 2015). **C**. TMD III residues (yellow fills) that influence the fraction of kiss-and-run events are illustrated in structure models from (Adams et al., 2015) (**C1**) and from Alphafold (**C2**) (Jumper et al., 2021).

We also used these structural models to interpret changes in release mode. M136, L139, and A142 tryptophan mutants increased the proportion of kiss-and-run events, and two of these, M136 and L139, increased the duration (Fig. 3). Fig. 7C1 highlights these residues in the electron microscopy structure. M136 is close to syb and L139 and A142 are close to syp TMD IV. However, in the Alphafold model, M136 and L139 face away from TMD IV and could potentially face toward syb (Fig. 7C2). Though the orientation of TMD III is different in the two models, both are consistent with the idea that kiss-and-run fusion pores are influenced by intermolecular interactions with syb and intramolecular interactions with other syp TMDs.

### Conclusions

TMD III of syp influences the dynamics of initial fusion pores without directly contacting the pathway taken by catecholamine as it leaves a vesicle. Although these results indicate that the syp-syb complex revealed by electron microscopy (Adams et al., 2015) is not likely to be the initial fusion pore, our results suggest an intimate relationship between syp and the initial fusion pore that enables TMD III to regulate fusion pore activity. Interactions between syp and its binding partner syb may play a role, but syp would then surround the pore-lining syb TMDs rather than alternate with them. The helical faces of syb with pore-lining residues (highlighted as purple fills in Figs. 7B1 and 7C1) are exposed to the pore lumen, inaccessible to contact with a syp TMD. Syp TMDs would then be more likely to interact with a face of syb that is oriented away from the pore lumen. A recent study showed that primed synaptic vesicles have a hexameric arrangement of protein density at the interface with the plasma membrane, surrounding the area where the initial fusion pore is likely to form (Radhakrishnan et al., 2021). Determining whether syp and syb contribute to this density, where they reside, and how they interact will suggest more experiments that can evaluate the functions of syp and syb in membrane fusion.

## Materials and Methods

### Animals

Breeding homozygous syp KO/heterozygous syg KO mice generated litters of ∼25% DKO (Eshkind and Leube, 1995; Janz et al., 1999). DKO offspring were identified by genotyping. All mice were housed and handled in accordance with the guidelines of the National Institutes of Health, and approved by the Animal Care and Use Committee of the University of Wisconsin-Madison.

### Chromaffin cell culture

Chromaffin cells were prepared as described previously (Chang et al., 2021). Mouse adrenal glands were collected in ice cold Hanks’ balanced salt solution (HBSS) with penicillin/streptomycin, cut into small pieces, and dissociated in DMEM with 20 units papain, 10 mg L-cysteine, 0.02 M CaCl_2_ and 0.02 M EDTA at 37°C for 25–30 min. After replacing the papain solution with enriched medium (DMEM with insulin-transferrin-selenium-X and penicillin/streptomycin), the tissue was triturated and plated on glass coverslips coated with poly-D-lysine for 1 h. One mL enriched medium was added and cells were maintained in an incubator at 37°C ventilated with a humidified air-5% CO_2_ atmosphere. Cells were used within 4 days.

### Molecular biology

Point mutations with tryptophan in TMD III were generated by modified sequential PCR (Cormack, 2008) with rat wild-type syp (pLox-CMV-syp-IRES2-EGFP) as the template. Primers were designed with the QuickChange Primer Design Program (Agilent Technologies). Constructs were confirmed by DNA sequencing, subcloned into a bicistronic lentiviral vector, pLox-CMV-IRES-EGFP, and purified. Lentivirus particles were generated in HEK 293T cells as described previously (Chang et al., 2021). DKO chromaffin cells were infected with virus particles on the day of dissection. Amperometry recordings were conducted 48-72 hours after viral infection.

### Amperometry

Chromaffin cells were recorded in bathing solution with 150 mM NaCl, 4.2 mM KCl, 1 mM NaH_2_PO_4_, 0.7 mM MgCl_2_, 2 mM CaCl_2_, and 10 mM HEPES (pH 7.4) at 22°C. Secretion was induced by pressure ejection of a high-KCl depolarizing solution consisting of 105 mM KCl and 2 mM NaCl, from a 2-μm tipped micropipette positioned 15-20 μm away. Catecholamine-oxidization currents were recorded with a 5-μm diameter-tipped carbon fiber electrode (CFE-2, ALA Scientific Instruments) polarized at 650 mV with a VA-10X amplifier (ALA Scientific Instruments). Signals were low-pass-filtered at 1 kHz and read into a computer at a digitization rate of 4 kHz with pClamp10 (Axon Instruments, Molecular Devices Corp). Depolarizing solution was applied 3 s after the start of acquisition for 6 s, and recording continued for another 14 s (total time, 23 s). Each cell was stimulated six times. Data analysis was conducted with an in-house computer program. Events with peak amplitude ≥ 2 pA were analyzed. Events 2-4 pA were taken as kiss-and-run, while events ≥ 4 pA were taken as full fusion (Chang et al., 2021). Spike frequency was calculated for events ≥ 4 pA from the first secretion episode of each cell. Rising-slopes were obtained by linear regression of the 10-70% rising phase of cumulative spike-count plots. Prespike feet (PSF) were analyzed from spikes ≥ 20 pA and mean PSF duration determined by fitting lifetime distributions with a single exponential decay. Spike characteristics and late-stage permeability were analyzed from spikes with peak amplitudes ≥ 20 pA using methods described previously (Jackson et al., 2020).

### Immunocytochemistry and confocal imaging

DKO chromaffin cells were co-transfected with WT or mutant syp together with neuropeptide Y-DsRed (NPY-DsRed). Cells were fixed 72 h after transfection with 4% paraformaldehyde for 30 min, permeabilized with 0.1% Triton X-100, and blocked in 10% goat serum for 1 h, all at room temperature (RT). Cells were then incubated at 4°C overnight with rabbit anti-syp antibody (Synaptic Systems #101002) and mouse anti-syb antibody (Synaptic Systems #104211) or mouse anti-RFP antibody (ThermoFisher #MA5-15257) at a dilution of 1:280. After washing three times with phosphate buffered saline (PBS) for 10 min, cells were incubated with AF568-goat anti-mouse (Abcam #ab175473) and AF647-goat anti-rabbit (Sigma #SAB4600185) antibodies, or AF647-goat anti-mouse (Sigma #SAB4600392) and AF546-goat anti-rabbit (Invitrogen #A11010) antibodies at 1:1000 dilution at RT for 1 h. Cells were then washed three times with PBS for 10 min, and incubated with DAPI at RT for 10 min. The coverslips were sealed onto slides with Fluoromount G (Electron Microscopy Sciences). Cells were imaged with a Leica TCS SP8 laser-scanning confocal microscope equipped with a 63x oil immersion lens. The laser intensity was set to 12% for 561 nm and 633 nm, and 5% for 405 nm. Pearson’s correlation coefficient was determined by Leica application suite X (LAS X) colocalization software.

### Statistics

All data were presented as mean ± SEM. Means of spike frequency, spike characteristics, PSF amplitude, fraction of kiss-and-run events, kiss-and-run duration, and permeability of fusion pores were reported as the cell-means, with averages first taken over each cell, and then averaging these means; cell number was used to calculate error (Colliver et al., 2000). One-way ANOVA followed by *post-hoc* Student-Newman-Keuls test, or the Kruskal-Wallis method followed by *post-hoc* Dunn test was used to evaluate statistical significance among different test groups. The significance was represented with asterisks with the following notation: * *p* < 0.05; ** *p* < 0.01; *** *p* < 0.001. All data and statistical analysis were performed with the computer programs Origin and GraphPad.

## Acknowledgements

We thank Che-Wei Chang for discussions and advice and Edwin Chapman for comments on the manuscript. This work was funded by NIH grant NS044057.

